# Immune landscape of isocitrate dehydrogenase stratified human gliomas

**DOI:** 10.1101/2022.11.08.514794

**Authors:** Pravesh Gupta, Minghao Dang, Shivangi Oberai, Mekenzie Peshoff, Nancy Milam, Aml Ahmed, Krishna Bojja, Tuan M. Tran, Kathryn Cox, Huma Shehwana, Carlos Kamiya-Matsuoka, Jianzhuo Li, Joy Gumin, Alicia Goldman, Sameer A. Seth, Atul Maheshwari, Frederick F. Lang, Nicholas E. Navin, Amy B. Heimberger, Karen Clise-Dwyer, Linghua Wang, Krishna P. Bhat

## Abstract

The brain tumor immune microenvironment (TIME) continuously evolves during glioma progression, but only a limited view of a highly complex glioma associated immune contexture across isocitrate dehydrogenase mutation (IDH) classified gliomas is known. Herein, we present an unprecedentedly comprehensive view of myeloid and lymphoid cell type diversity based on our single cell RNA sequencing and spectral cytometry-based interrogation of tumor-associated leukocytes from fifty-five IDH stratified primary and recurrent human gliomas and three non-glioma brains. Our analyses revealed twenty-two myeloid and lymphoid cell types within and across glioma subtypes. Glioma severity correlated with microglial attrition concomitant with a continuum of invading monocyte-derived microglia-like and macrophages amongst other infiltrating conventional T and NK lymphocytes and unconventional mucosa associated invariant T (MAIT) cells. Specifically, certain microglial and monocyte-derived subpopulations were associated with antigen presentation gene modules, akin to cross-presenting dendritic cells (DCs). Furthermore, we identified phagocytosis and antigen presentation gene modules enriched in Triggering receptor expressed on myeloid (TREM)-2^+^ cells as a putative anti-glioma axis. Accelerated glioma growth was observed in *Trem2* deficient mice implanted with CT2A glioma cells affirming the anti-glioma role of TREM2^+^ myeloid cells. In addition to providing a comprehensive landscape of glioma-specific immune contexture, our investigations discover TREM2 as a novel immunotherapy target for brain malignancies.

## INTRODUCTION

The discovery of meningeal and parenchymal access of immune cells^1^ and the presence of meningeal^2-4^ and dural lymphatics^5^ in the central nervous system (CNS) has led to redefinition of the brain being immunologically distinct rather than immune privileged; a notion held for several decades^6^. In brain pathologies such as traumatic injury or neurodegenerative disorders, phagocytic cells comprising brain resident microglia and CNS associated monocytes/macrophages are the first responders to resolve associated inflammation. For instance, in Alzheimer’s disease the protective role of microglia in clearing amyloid plaques has been established to prevent disease progression^7^. In contrast, tumor associated macrophages have been linked to poor prognosis in brain neoplasms such as gliomas that arise from transformed neural progenitor cells^8,9^. Although primarily phagocytic in nature, myeloid cells are plastic and can undergo functional diversification under the influence of dysregulated cytokine and chemokine milieu contributed by both infiltrating bone marrow derived leukocytes as well as tumor cells^10-13^. Myeloid cell functions can also be differentially influenced by tumor necrosis and inflammation, a defining feature of IDH-wt when compared to IDH-mut gliomas^14^. Standard of care treatments such as surgical resection followed by temozolamide and ionizing radiation can unintendedly cause disruption of anatomical barriers, immunomodulation and necrosis, all of which can skew the properties of myeloid cells and other leukocytes^15,16^.

Studies pertaining to immune cell heterogeneity of gliomas at single cell resolution are emerging, however these studies are either restricted to myeloid cells^13,17-21^ or lack in-depth characterization of low grade in comparison to high grade gliomas^17,18,22-24^. Furthermore, given that most immunotherapy clinical trials are prioritized in relapsed patients^25^, understanding the treatment induced changes of brain TIME with unbiased approaches in recurrent tumors of all glioma subtypes are imperative. Glioma specific TIME studies are broadly focused on microglia /macrophages collectively referred as glioma-associated macrophages (GAMs), myeloid-derived suppressor cells (MDSCs), and tumor infiltrating lymphocytes (TILs)^23,26^. However, oversimplified myeloid cell diversity and M1/M2 functional dichotomization ignores the phenotypic heterogeneity and plasticity in these cell types^27-29^. Recent cytometry studies have captured the leukocyte diversity of the brain TIME in primary gliomas and brain metastasis^30,31^ to a certain extent albeit based on *a priori* markers.

To address these knowledge gaps and delineate the glioma associated leukocyte diversity in the TIME, we performed single cell (sc)- and bulk RNA Sequencing (RNA-seq) and spectral cytometry analyses on tumor-associated leukocytes from fifty-five IDH-stratified primary (treatment naïve) and standard of care treated recurrent glioma subtypes to define their immune landscape. In addition to corroborating established myeloid dominant glioma characteristics, we redefine glioma TIME by superimposing our advanced findings across glioma subtypes with largely IDH-wt glioma restrictive studies^17,18,22-24^. Our major findings include the following: i) We observed significant attrition of microglia (MG) accompanied by increased infiltration of classical monocytes (c-Mo), monocyte-derived-microglia-like (Mo-MG or MG-like), -macrophages (MDM) and conventional dendritic cells (cDC)-2 in recurrent IDH-wild type gliomas relative to other glioma subtypes; ii) We demonstrate eleven transcriptionally distinct glioma associated MG states inclusive of tumoricidal, inflammatory and metabolic phenotypes; ii) Infiltration of Tregs, NK cells and mucosa associated invariant T (MAIT) were significantly abundant in recurrent IDH-wild type gliomas; and iv) We identified glioma associated myeloid cells with triggering receptor expressed on myeloid (TREM)-2 cells enriched for phagocytosis and antigen-presentation gene modules as putative anti-glioma axis and demonstrate their anti-glioma functions using a xenograft mouse model. In summary, our reverse translational glioma immunophenotyping investigations reveal an unprecedently advanced landscape of glioma TIME that can be exploited for future immunotherapy applications. We further uncover TREM2 as a novel glioma specific immunomodulatory target with likely implications in other brain malignancies.

## RESULTS

### Transcriptionally defined immune cell diversity in IDH-mutation stratified human gliomas

To discern glioma associated immune cell diversity, we performed single cell RNA sequencing (scRNA-seq) on flow sorted CD45^+^ leukocytes obtained from tumors of eighteen IDH-mutation classified patients comprising IDH-mutant primary (IMP; n=4), IDH-mutant recurrent (IMR; n=6), IDH-wild type primary (IWP; n=4), or IDH-wild type recurrent (IWR; n=4) gliomas (hereafter referred as glioma subtypes). Three quasi-normal, non-glioma brains (NGBs) either from a grade I meningioma patient or refractory epileptic non-neoplastic patients were used as controls (**Fig. 1A and Supplementary Table S1**). Using a previously described immune cell enrichment protocol^32^ and CD45^+^ sorting strategy, we consistently obtained highly pure CD45^hi^/CD45^lo^ leukocyte subpopulations in gliomas and NGBs (**Fig. 1A and Supplementary Fig. S1A-C**). This is in contrast to previous studies using human glioma specimens that were not able to resolve these two distinct CD45^hi^/CD45^lo^ subpopulations^33-35^.

**Fig. 1.**
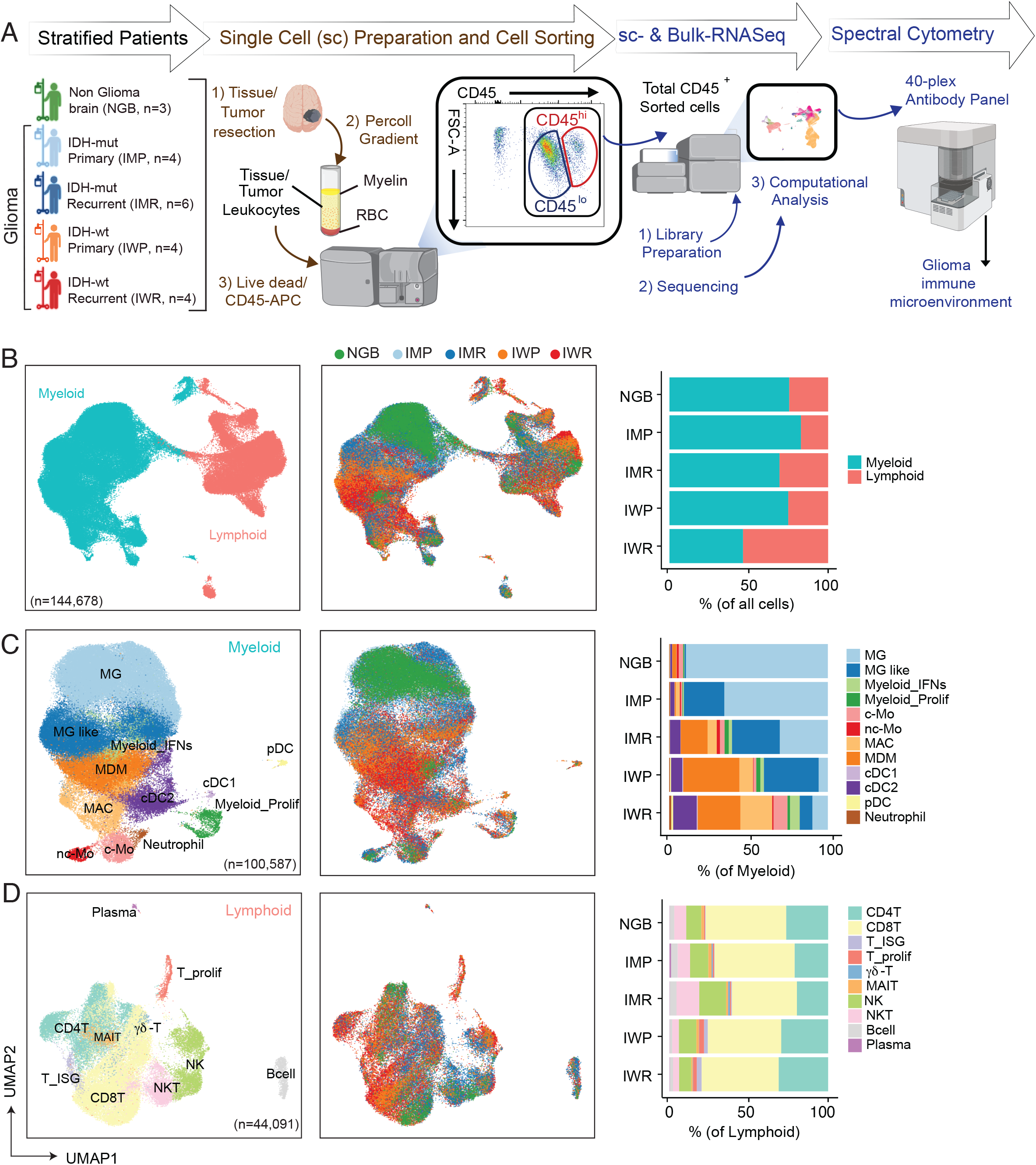
The single cell transcriptional landscape of glioma TIME. **A)** A schema depicting the experimental workflow from sample preparation (see Methods) of resected brain tissues/tumors to scRNA-seq data generation and spectral flow cytometry validation and computational analysis. Patients were stratified as non-glioma brain (NGB; n = 3) and IDH-mutant primary (IMP; n = 4), IDH-mutant recurrent (IMR; n = 6), IDH-wild type primary (IWP; n = 4), IDH-wild type recurrent (IWR; n = 6) groups; hereafter collectively referred as glioma subtypes (see details in **Supplementary Table 1**). The dissociated CD45-APC-stained cells were FACS sorted to obtain pure CD45^+^ glioma associated leukocytes. Subsequently matched sc-RNAseq and bulk RNA-Seq was performed followed by computational analysis. sc-RNA seq inferred cell types were validated by spectral cytometry. **B-D)**, Uniform manifold approximation and projection (UMAP) visualization of unsupervised clustering analysis of **(B)** all immune cells (n = 144,678) that passed quality filtering (see Methods), **(C)** myeloid lineage clusters (n = 100,587), and **(D)** lymphoid lineage clusters (n = 44,091). Cells are color coded for their inferred cell types (left) and the glioma subtypes of their corresponding tumors (middle).

Overall scRNA-seq dataset was batch corrected with Harmony using COMBAT software **(Supplementary Fig. S1D-G)**. A low rejection rate in K-BET indicated homogeneous mixing of samples after batch correction (**Supplementary Fig. S1H**) as detailed in methods. Furthermore, we removed the likely doublets/multiplets, cell debris, and low-quality cells using a multistep approach (see Methods). Unsupervised clustering followed by Uniform Manifold Approximation and Projection (UMAP) analyses resolved 144,678 cells into two major immune compartments predominated by myeloid (n=100,587) and lymphoid cells (n=44,091) within and across glioma subtypes in line with previous reports^30,31,36,16,17,21-23^ with the exception of IWR gliomas, which had similar proportions of myeloid and lymphoid cell populations (**Fig. 1B)**. Glioma associated myeloid cells exhibited a continuum of overlapping gene features (*SPP1, APOE, C1QC*) with MG and macrophage (MAC) subsets yet resolved by core MG gene set including *CX3CR1, GPR34, P2RY12, P2RY13, SALL1, TAL1*, and *TMEM119* amongst others (**Fig. 1C, Supplementary Fig. S1J)**. Dendritic cells (DC), monocytes (Mo), and neutrophils were clearly identified based on expression of canonical gene signatures (**Fig. 1C, Supplementary Fig. S1I and J)**.

Brain resident MG represented the largest myeloid cell type in NGB and IMP (**Fig. 1C**). In contrast, in response to MG attrition, invading MG-like cells co-expressing MG- (*SORL1, SAMD9L, GPR34*) and MAC- (*GLDN, MSR1, CD163*) signature genes, *VCAN*^*+*^*FCN1*^*+*^ classical monocytes (c-Mo), *TCF7L2*^*+*^*FCGR3A*^*+*^ non-classical monocytes (nc-Mo), *CD163*^*+*^*MARCO*^*+*^*FN1*^*+*^ MAC, *TMEM176A*^*+*^*SELENOP*^*+*^ MDM proportionately increased in IMR, IWP and IWR glioma subtype (**Fig. 1C, Supplementary Fig. S1J**). Increased trends with professional antigen presenting cells (APC) such as *CLEC9A*^*+*^ cDC1, *CD1C*^*+*^ cDC2 and *IL3RA*^*+*^ plasmacytoid DC (pDC) were observed in IWR gliomas compared to other glioma subtypes. Other notable glioma associated myeloid cells included indistinguishable cell type with enriched interferon (IFN) stimulated gene signatures (*IFI44L, IFI6, ISG15*) defined as MAC/MG_IFNs, and proliferative genes (*MKI67, PCLAF*) expressing Myeloid_Proliferative (Myeloid_prolif) cells and *JMJD1C*^+^ Neutrophils (**Fig. 1C, Supplementary Fig. S1J**). We speculate that the influx of non-MG myeloid cells is a consequence of depleting niches of MG akin to observations in inflammation associated conditions where tissue resident macrophage attrition has been reported^37^.

Inter- and intratumoral glioma associated lymphoid cell types resolved into T lymphocytes, *TRDC*^*+*^ γδ-T cells, *SLC4A10*^*+*^ Mucosa associated invariant T (MAIT) cells, *NKG7*^*+*^ *KLRF1*^*+*^ Natural Killer (NK) cells, *CD3D*^*+*^*NKG7*^*+*^ Natural Killer T (NKT) cells, *CD79A*^*+*^*MS4A1*^*+*^B lymphocytes and *MZB1*^*+*^*IGHG1*^*+*^ plasma cells (**Fig. 1D, Supplementary Fig. S1K and 1L)**. Notably, amongst lymphoid lineage cells, we identified rare infiltrating populations of MAIT (0.18%-4.14%) and NKT (0.35% - 26.4%) in human glioma TIME, which have not been previously reported^38^. We observed glioma subtype specific enrichment patterns, such as apparently reduced MG and abundance of T-cell and monocyte-derived cells in all patients in the IWR group **(Supplementary Fig. S1M)**. Overall, we report twelve myeloid and ten lymphoid cell types with transcriptionally defined phylogenic cellular relationships in the TIME of IDH stratified primary and recurrent human gliomas **(Supplementary Fig. S1N)**.

### Glioma associated transcriptional immune cell phenotypes validated by spectral cytometry

In view of transcriptionally redefined immune cell diversity across glioma subtypes, we validated scRNA-seq inferred cell types with correlative protein markers. A 40-parameter protein marker panel was designed (**Supplementary Table S2**) to corroborate majority of inferred cell types **(Supplementary Fig. 1N)**. A comprehensive spectral cytometry-based phenotyping immunoassay was performed across fifty-five patients covering all glioma subtypes and three refractory epileptic non-neoplastic patients (detailed in **Supplementary Table S1**). We confirmed P2RY12^+^CX3CR1^+^MG as the most abundant cells across glioma subtypes with highest proportions evident in NGB and IMP (**Fig. 2A**). Total P2RY12^+^CX3CR1^+^MG and even reactive CD11c^+^MG were dramatically reduced with glioma recurrence in IMR and IWR compared to IMP and IWP glioma subtype respectively. A concomitant significant increase in CD11c^+^CCR2^+^Mo-MG, CD14^+^CD16^-^c-Mo, CD68^+^CCR2^+^MDM and CD1c^+^cDC2 was observed in IWR relative to IMP glioma subtype while other myeloid cell types such as CD16^-^CD14^+^nc-Mo, CD68^+^CCR2^+^MDM, Clec9A^+^cDC1 (**Fig. 2 B-E**) and CD66b^+^ neutrophils (not shown). Amongst lymphoid lineage cells Foxp3^+^Tregs, CD56^hi^NK, CD56^lo^NK cells and TCRVα7.2^+^mucosa associated invariant T (MAIT) were significantly abundant in IWR gliomas with co-presence of CD4, CD8 T, NKT, and ψο-T cells across glioma subtypes (**Fig 2F-H**). Altogether, we confirmed all major cell types and their enrichment patterns across glioma subtypes.

**Fig. 2.**
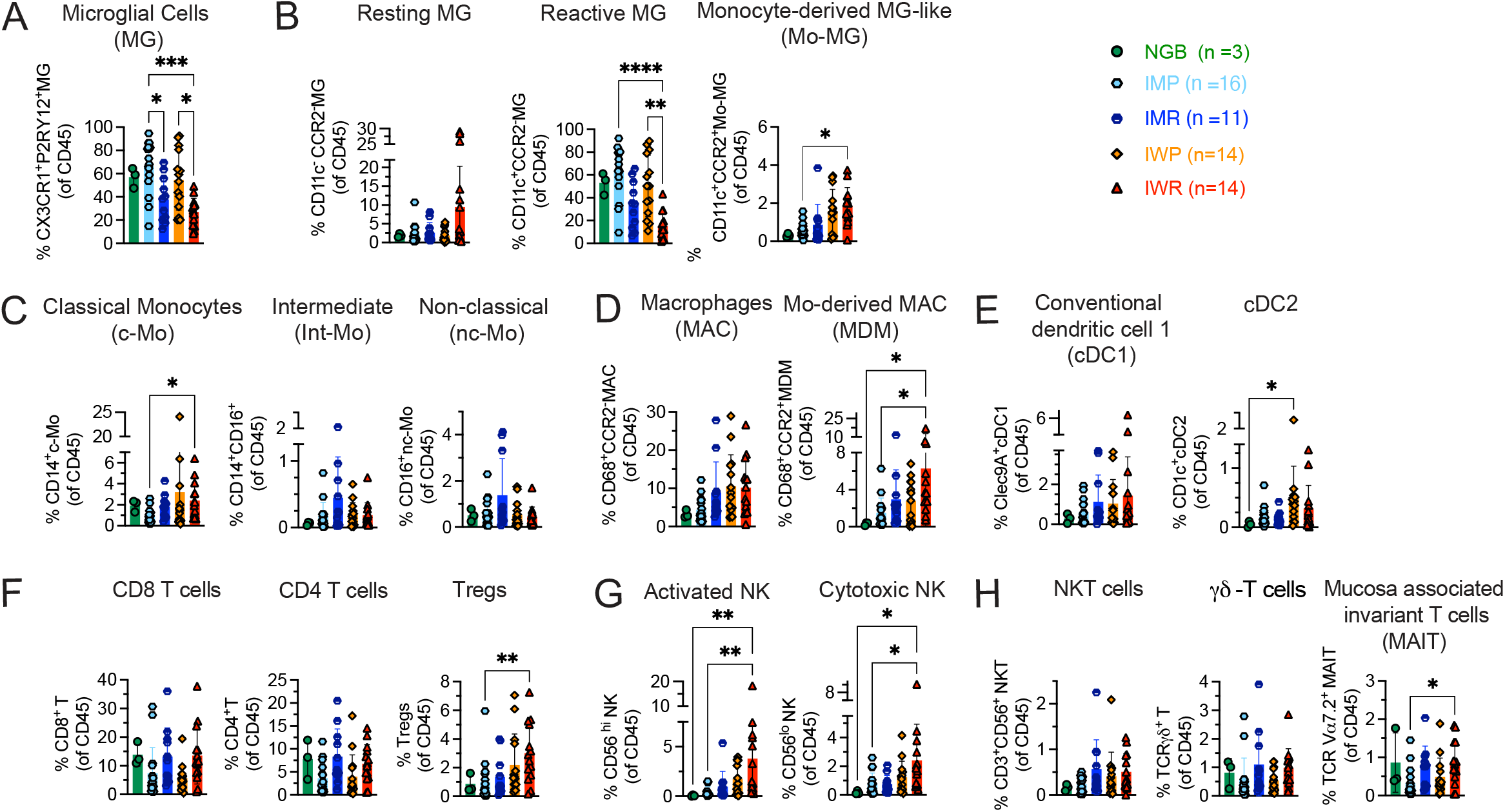
Glioma associated leukocyte diversity corroborated with spectral flow cytometric analyses. Scatter bar plots represent the proportion of indicated immune cell type (of total CD45^+^ leukocytes) across glioma subtypes; NGB (n = 3), IMP (n = 16), IMR (n = 11), IWP (n = 14) and IWR (n =14) as shown in **(A)** CX3CR1^+^P2RY12^+^ Microglial cells (MG); **(B)** Microglial subsets (gated on MG): CD11c^-^CCR2^-^ Resting MG (left), CD11c^+^CCR2^-^ Reactive Microglia (middle), and CD11c^+^CCR2^+^ Monocyte-derived MG-like (Mo-MG); **(C)** Monocyte subsets (gated on CD3^-^ CD56^-^): CD14^+^CD16^-^ Classical Monocytes (c-Mo), and CD14^+^CD16^+^ Intermediate monocytes (Int-Mo) and CD14^-^CD16^+^ Non-classical monocytes; **(D)** Macrophages (Gated on CD3^-^CD56^-^): CD68^+^CCR2^-^ Macrophages (MAC), and CD68^+^CCR2^+^ Monocyte-derived Macrophages (MDM) **(E)** Conventional Dendritic cell (cDC) subsets (gated on CD11b^+^CD11c^+^HLADR^+^): Clec9A^+^CD1c^-^ cDC1, and Clec9A^-^CD1c^+^ cDC2; **(F)** T cell subsets (Gated on CD3^+^); CD8^+^CD4^-^ T cells, CD4^+^Foxp3^-^ T cells and CD4^+^Foxp3^-^ T regulatory (Tregs) cells; **(G)** Natural Killer cell subsets (gated on CD56^+^); CD56^hi^CD16^lo^ Activated NK cells, and CD56^lo^CD16^hi^ Cytotoxic NK cells; **(H)** Unconventional T cell subsets: CD3^+^CD56^+^ Natural Killer T (NKT) cells, TCRψο^+^ (ψο-T) cells TCRVα7.2^+^ Mucosal associated invariant T (MAIT) cells. Statistical significance was determined using by Kruskal Wallis test at p*<0.05, p**<0.01, p***<0.001.

### Transcriptional heterogeneity and inferred functional states of microglial cells in human gliomas

In humans, the CNS associated microglial states have been defined in Alzheimer’s^39^, multiple sclerosis^40^ and to a limited extent in primary IDH-wt and IDH-mut gliomas using single cell transcriptomics^17,18,23,41^. We sub-clustered and delineated MG into eleven distinct states distributed across glioma subtypes (**Fig. 3A, Supplementary Fig. S2B-E)**. These MG clusters were distinguished from the rest of the myeloid cells based on the core MG gene sets inclusive of MG specific transcription factors; *TAL1* and *SALL1* **(Fig. S1J)**.

**Fig. 3.**
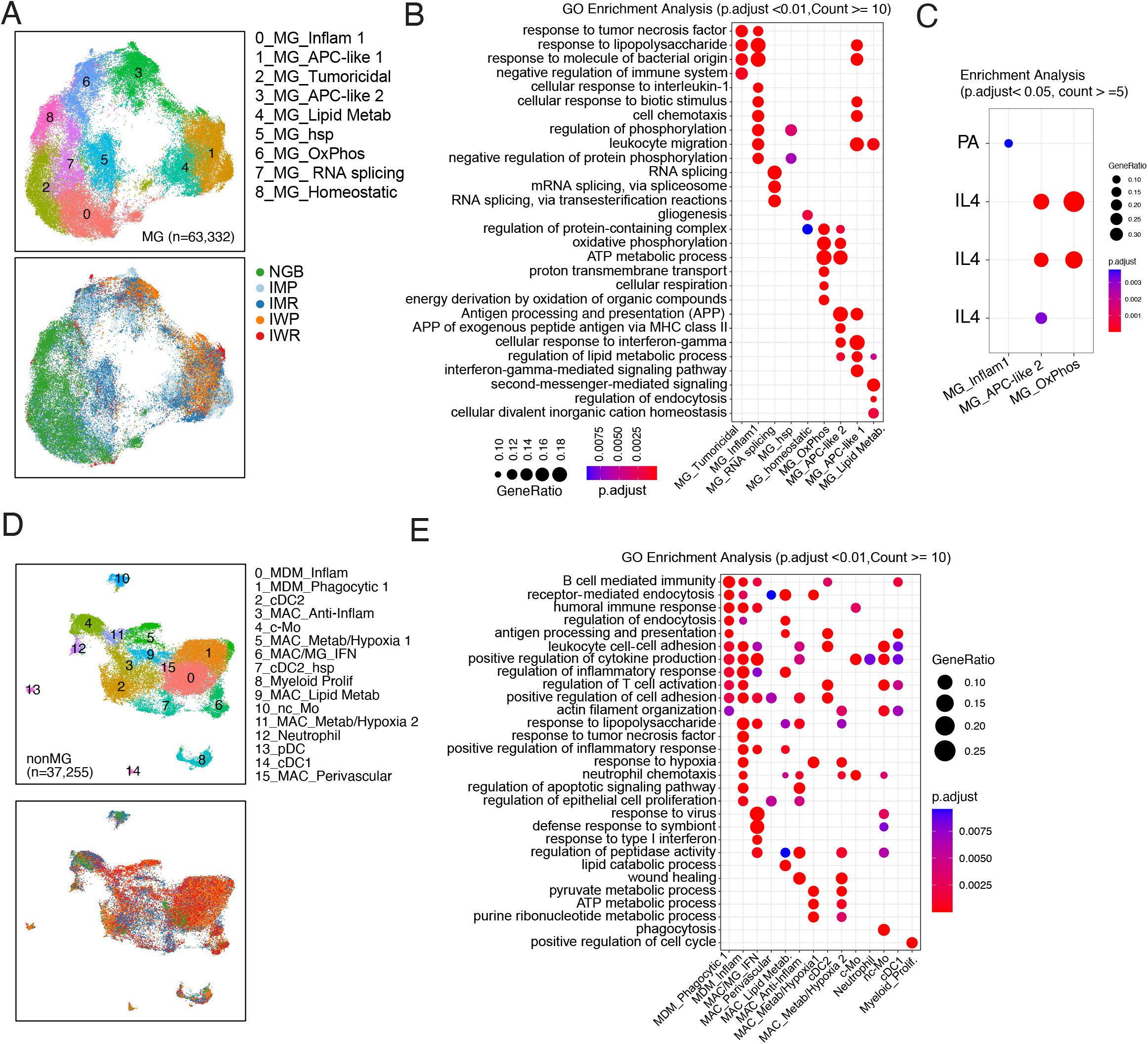
Glioma associated myeloid cell diversity and their inferred functional states. **(A)** UMAP visualization of unsupervised clustering analysis of Microglia (MG, n = 63,332), displaying nine distinct cell states (top). Cells are color coded for their inferred cell types (top) and the glioma subtypes of their corresponding tumors (bottom). **(B)** Bubble plot depicting gene ontology (GO) analysis of glioma associated MG states. **(C)** Bubble plot showing overrepresented stimulus-specific Palmitic acid and Interleukin-4 (IL-4) polarization gene expression modules as multispectral polarization (see Methods). **(D)** UMAP visualization of unsupervised clustering analysis of non-MG myeloid cells (n = 37,255), displaying sixteen distinct cell states (top). Cells are color coded for their inferred cell types (top) and the glioma subtypes of their corresponding tumors (bottom). **(E)** Bubble plot depicting gene GO analysis of glioma associated non-MG myeloid cell states. **(B, C, E)** In B and E, each bubble represents a GO Term and in C, each bubble represents a polarization module. Bubble size corresponds to gene ratio and the color of the bubble indicates statistical significance. Only polarization modules with 5 or more overlapping genes and an adjusted p value of <0.05 are shown in C.

To evaluate functional features, we performed gene ontology (GO) analysis which identified MG expressing unique markers with subtype specific clustering. This included glioma restricted antigen presentation associated gene modules (e.g., *CSTD, MS4A4A)* in MG_APC-Like 1 and MG_APC-like 2 clusters and a lipid metabolism associated *LPL*^*+*^ MG_Lipid Metab. Cluster **(Fig. 3B, Supplementary Fig. S2C)**. We also observed IDH-mut glioma restricted MG clusters that included *BAG3*^+^MG_hsp and *ATF* expressing metabolically enriched MG_OxPhos **(Fig. 3B, Supplementary Fig. S2C)**. MG that was predominantly seen in NGB were *P2RY12*^*+*^ MG_homeostatic and *CCL4L2* expressing inflammatory MG_Inflam 1 cluster associated with response to tumor necrosis factor, Interleukin-1 and lipopolysaccharide GO term **(Fig. 3B, Supplementary Fig. S2C)**. Tissue macrophages exhibit multifaceted polarization in response to microenvironmental cues, hence we evaluated the polarization spectrum of MG with nine distinct macrophage activation programs as previously described^27^. We assessed polarization states of MG clusters with a spectral polarization view rather than dichotomous M1/M2 polarization model and identified palmitic acid responsive gene module associated with MG_Inflam1 along with IL-4 responsive polarization states in MG_APC-like 2 and MG_OxPhos clusters **(Fig. 3C)**. A *GNLY*^*+*^*TNF*^*+*^ cluster defined as MG_Tumoricidal was also noted across glioma subtypes. Furthermore, we subclustered the unidentifiable MAC/MG_IFNs cluster, which enabled resolution of this cluster into MAC and MG phenotypes. These *IFN* gene associated clusters were *CD163*^*+*^*LYZ*^*+*^MAC_IFN, *GNLY*^*+*^*TMIGD3*^*+*^MG_IFN *GNLY*^*+*^*IL1A*^*+*^MG Inflam 2 **(Fig. 3C, Supplementary Fig. S2D-E)** and associated gene signatures are suggestive of their contribution to inflammation in gliomas. Overall, we identified transcriptionally heterogeneous MG with their inherent functional likelihoods across glioma subtypes.

### Spectrum of invading non-MG myeloid cells in human gliomas

In order to understand non-MG myeloid cell diversity, which have been a subject of intense investigation in IDH-wt gliomas^12,13,18,20,21,33^, we performed a comprehensive analyses of invading non-MG myeloid subpopulations across IDH-classified gliomas and identified sixteen non-myeloid cell states **(Fig. 3D)**. We uncovered six MAC and two MDM clusters. Based on DEG and GO analysis we annotated glioma associated macrophages as *IL10*^*+*^MAC_Anti-Inflam, multiple metabolic phenotypes such as clusters enriched with hypoxia genes (e.g., *SDS, HMOX1*) MAC_Metab/Hypoxia 1, MAC_Metab/Hypoxia 2 and *LIPA*^*+*^MAC_Lipid Metab subpopulations. A cluster of *LYVE1* expressing MAC associated with vasculature defined as MAC_Perivascular was observed across glioma subtypes **(Fig. 3E, Supplementary Fig. S2F-G)**. Amongst the MDMs, *C1QA*^*+*^MDM_Phagocytic 1 was associated with receptor mediated endocytic process and *JUN*^*+*^*SELENOP*^*+*^MDM_Inflam subset correlated with positive regulation of inflammation GO term **(Fig. 3E, Supplementary Fig. S1M, S2F-G)**. Although MDMs showed marked increase in IWR glioma subtype, abundance of MAC was proportionately similar across IMP, IMR and IWP gliomas (**Fig. 2D**). Monocytic infiltrates in glioma TIME included significantly abundant *FCN1*^*+*^*CD14*^*+*^c-Mo in IWR compared to IMP gliomas whereas *FCGR3A*^+^*TCF7L2*^+^ nc-Mo did not show any noticeable differences across other glioma subtypes (**Fig. 3D, 2C, Supplementary Fig. S1M, S2F**). Glioma associated inflammation led to DC infiltration in contrast to negligible DCs in NGB *CD1C*^*+*^cDC2 was significantly abundant in IWR glioma subtypes, while a proportionally similar levels of infiltration of *CLEC9A*^+^cDC1 and *IL3RA*^+^pDC were observed across other glioma subtypes (**Fig. 3D, 2E, Supplementary Fig. S1M, S2F-G**). Taken together, our data provides advanced insight of infiltrating myeloid cell subsets in the glioma TIME and highlights their similarities and differences with MG.

### Identification of Trem2 as an anti-glioma modulator

To gain a deeper understanding of pathways altered in MG, we inspected genes that regulate phagocytosis and antigen presentation in myeloid cells, especially MG_APC-Like 1 and MG_APC-like 2 clusters (**Fig. 4A**). These clusters were enriched for *HLA-DR* (**Fig. 4A**), a major histocompatibility complex (MHC) class II gene. In addition, we found higher expression of *TREM2* and its regulator *MS4A6A* (**Fig. 4B**), both of which play crucial roles in microglial functions and gene risk loci for Alzheimer’s disease, where MG play a protective role. Recent studies have also demonstrated a role for Trem2 in immunosuppression of cancer ^42,43^. To examine the influence of TREM2 on glioma growth, we implanted CT-2A glioma cells in C57BL/6 WT mice and compared survival differences with Trem2^-/-^ mice. Genetic ablation of Trem2 promoted tumor growth in mice (bioluminescence data not shown) and showed significantly reduced survival compared to syngeneic WT mice with intracranial gliomas (**Fig. 4C**). These data demonstrate that in contrast to previous studies on systemic cancers^44,45^, TREM2^+^ myeloid cells play an anti-tumor role in GBM.

**Fig. 4.**
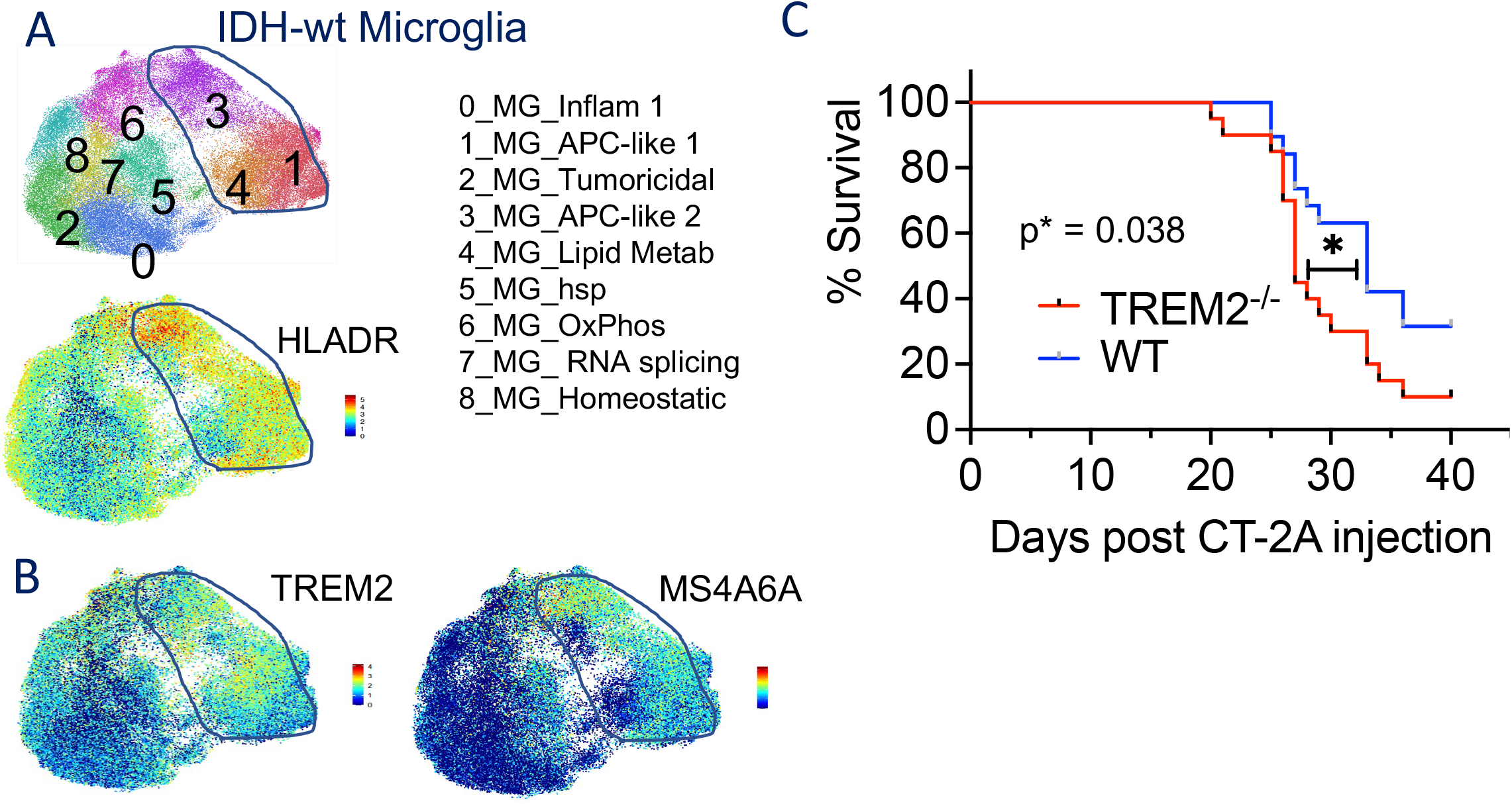
Identification of anti-glioma role of Triggering receptor expressed on myeloid (TREM)-2^+^ myeloid cells. **(A)** UMAP visualization of (top) inferred MG states, expression of *HLADR* (bottom); **(B)** TREM2; (**C)** MS4A6A across MG clusters as shown in (A). **(D)** Representative percentage survival analyses of CT-2A glioma bearing mice TREM2^-/-^ (n=20) versus WT mice (n=19) in C57BL/6 genetic background. Statistical significance of survival was determined using Mantel-Cox log-rank test at p*=0.038.

## DISCUSSION

Using single cell transcriptomic profiling, our study uncovers the cellular and molecular landscape of brain-specific glioma immunity. We observed that both myeloid and lymphoid subpopulations exhibit remarkable cellular diversity depending on IDH status and disease severity. Recent high throughput studies have elegantly demonstrated the spatio-temporal distribution of microglial subsets in humans and mice^46^. These studies established the plasticity of microglial cell states and reveal mechanisms by which MG contribute to limiting or promoting neurodegenerative diseases^46^. We observed attrition of tissue-resident microglial cells with a concomitant increased infiltration of non-MG myeloid cells as a distinct feature in IDH-wt gliomas, which is consistent with recent reports^30,31^. In our investigation, we even report reduction in MG in relapsed IDH-mutant gliomas and the highest MG attrition evident in IWR gliomas. Acute inflammation induces transient loss of embryonically derived tissue-resident macrophages as a result of necroptosis and concomitant replenishment either through self-renewal or monocytic input as has been described in murine spleen, liver and lungs^37^. In line with this, we speculate migration of bone marrow derived myeloid cells as a compensatory mechanism of homeostatic myelopoiesis to fill depleting brain macrophage niches. Our results showing increased proportion of Mo-MG like cells in response to dramatic reduced MG in IWR gliomas provides a likely clue for such replenishment patterns in human gliomas. Our findings with MG provide evidence for multifaceted inflammatory phenotypes characterized by *IL1A, TNF, IL6, IL10 and GNLY* expression on various MG subsets. Although recent bulk mRNA-seq analysis pointed to cumulative spectral nature of glioma associated MG^31^, we clarify heterogeneity of polarization states and identify an unreported palmitic acid (PA) responsive and widely acknowledged IL-4 responsive gene modules in distinct clusters of MG. A Glucocorticoid induced signature has been reported with SEPP1^high^ Mo-TAMS^13^. Together our findings suggest distinct polarization states amongst the multispectral polarization background in glioma associated MG rather than M1/M2 dichotomy.

We observed that certain MG subsets abundantly expressed genes involved in both MHC class I (*B2M*) and class II (*HLA-DRB1*) molecules suggestive of their APC-like characteristics. With virtues of antigen presenting molecules, APC-like MG can potentially direct proliferation and secretion of cytokines in CD4 and CD8 T cells, a hypothesis that is worthy of further investigation. Interestingly we found that while MG express both MHC class I and class II gene modules; MAC or MDM are largely restricted to MHC class II. Although MG can orchestrate APC like functions, their reduced numbers may likely contribute to tumor immune escape. Recent studies have shown that MG localization is confined to the tumor border compared to macrophages which are enriched in the tumor core, implying that MG may have distinct spatio-phenotypic features compared to peripheral myeloid cells^22^. Our study also identified novel clusters of mo-DCs and MDMs that can also possess APC-like properties. Therefore, the dogma of DCs as *bona fide* immune sentinels and microglia and GAMs as being mainly tumor supportive needs to be revisited.

Despite the evidence of lymphopenia in glioma patients^47^ and minimal responsiveness or treatment refractoriness to checkpoint blockade interventions (e.g., nivolumab)^48^, GBM immunotherapy clinical trials are T cell centric. In this study, we report paucity of lymphocytes in primary gliomas, and therefore exclusive T cell-based therapies, may not be effective for such patient cohorts. In contrast, myeloid cells represented 50-80% of leukocytes compared to lymphocytes across gliomas. Recent studies have shown that targeting the innate immunity using either depletion strategies or inducing tumor phagocytosis as alternative immunomodulatory therapies in GBM^49^. Here, we show that TREM2 as a putative target with anti-glioma properties. Although TREM2 was shown to be immunosuppressive in other cancers, TREM2 genetic ablation induced glioma growth in mice indicating differential functions of TREM2 in brain tumors. Further investigation of TREM2 expression in brain resident and infiltrating immune cells is warranted.

In summary, our unbiased high dimensional studies have paved the way for an advanced understanding of immune landscape across gliomas that can be exploited for novel immunotherapy strategies for these cancer types.

## ACKNOWLEDGEMENTS

This study in K.B. laboratory was supported by the generous philanthropic contributions to The University of Texas (UT) MD Anderson Cancer Center (MDACC) Moon Shots Program™, Marnie Rose Foundation, NIH grants: R21 CA222992 and R01CA225963. This study was partly supported by the UT MDACC start-up research fund to L.W. and CPRIT Single Core grant RP180684 to N. E. N. We are thankful to Cynthia Kassab and Martina Ott for their contribution to tissue logistics for scRNA-seq pipeline. In part, the patient specimens analyzed with spectral cytometry in this study were contributed by CNS tumor Analysis Stream (CATALYST) program at UT MDACC and we extend our gratitude to entire CATALYST team. We thank Sanaalarab Al-Enazy for schematic illustrations. We would like to thank Haidee D. Chancoco for her contribution to RNA isolation for bulk mRNA-seq at Biospecimen Extraction Resource of MD Anderson Cancer Center; Advanced Technology Genomics Core supported by NCI grant CA016672 for library preparation and sequencing; and David W. Dwyer and Karen C. Dwyer at the Advanced Cytometry & Sorting Core Facility supported by NCI P30 CA016672.

## AUTHOR CONTRIBUTIONS

P.G. and K.P.B. conceived and designed the study. P.G., S. O., M.P., N.M., A.A. J.G., and K.B. acquired and processed glioma specimens. A.G., S.A.S and A.M. acquired epileptic clinical specimens. C.K.M. assisted in accessing the glioma patients’ clinical information. P.G., K.B., performed sample preparation for scRNA-seq. P.G. performed FACS and related data analysis. T.M.T. and J.L. performed scRNA-seq experiments. K.C.D. and PG designed cytometry panel. P.G, S.O. performed and analyzed spectral cytometry experiments under guidance of KCD. K.C. S.O. and P.G. acquired the data under guidance of K.C.D. Animal studies were performed by M.P. M.D. and H.S. performed bioinformatics analysis under the supervision of L.W and inputs from PG, and K.P.B. F.F.L., N.E.N. and A.B.H. provided constructive comments. P.G., M.D., L.W. and K.P.B. interpreted the scRNA-seq data. P.G., K.C.D. and K.P.B. interpreted the spectral cytometry data. P.G., M.D. L.W. and K.P.B. wrote and proofread the manuscript with contributions from all other authors.

## Data and Code Availability

All data and code available upon request

## METHODS

### Human brain tumor and tissue collection

Th**e** brain tumor/tissue samples were collected from seventeen-patients post appropriate informed consent during neurosurgery with detailed information pertaining to gender, age, glioma grade, subtype and, brain site of tumor extraction etc. mentioned in **Supplementary Table S1**. The brain tumor/tissue samples were collected as per MD Anderson internal review board (IRB)-approved protocol numbers LAB03-0687, LAB04-0001 and 2012-0441. Non-tumor brain tissue sample was collected from patient undergoing neurosurgery for epilepsy as per Baylor College of Medicine IRB-approved protocol number H-13798. All experiments were compliant with the review board of MD Anderson Cancer Center, USA.

### Preparation of leukocyte single cell suspensions from brain tumor and tissue

The resected brain tumors and, tissues were either freshly processed or transiently stored overnight MACS Tissue Storage solution (Cat. #130-100-008, Miltenyi Biotec) at 4 degree Celsius (in case of delayed surgeries) and processed immediately next morning. The brain tumor/tissue were finely minced and, enzymatic dissociation was performed in prewarmed digestion medium containing 100 μg/ml Collagenase D (Cat. #11088866001, Sigma-Aldrich) and, 2U/ml DNase (Cat. #D9905K/ NC0893386, Fisher Scientific) for 45 minutes at 37 degree Celsius. The enzymatic reaction was neutralized using 2% serum (Cat. #16140-071, gibco) in IMDM. The enzyme-digested tissues were homogenized by passing through an 18.5G gauge needle (Cat. #305196, BD) five times followed by further homogenization using syringe piston on a 100-μ cell strainer (Cat. #0877119/ Corning 352360) placed on a 50ml Falcon. The residual tissue on the strainer were mechanically dissociated using the piston of a 3ml syringe (Cat. #309657, BD). The single cells thus obtained were washed in 2% PBS and, centrifuged at 350g for 5 minutes at 4 degree Celsius. The resulting pellet was further subjected to 33% Percoll™ (Cat. #17-0891-01, Sigma-Aldrich) gradient and centrifuged at 800g without brakes for 12 minutes at 4 degree Celsius. The resulting pellet was subjected to the RBC lysis reaction (Cat. #R7757-100ML, Sigma-Aldrich) for 10 minutes at room temperature (R.T.) and reaction was stopped with 1X PBS (Cat. #21-040-CV, Corning) and resulting cell pellet obtained by centrifugation (500g, 5min, 4 degree Celsius). The single cells were filtered and cryopreserved in 10% DMSO (Cat. #D8418, Sigma-Aldrich) in FBS (Cat. #F4135, Sigma-Aldrich) in liquid nitrogen (−196 degree Celsius) after a transient storage at -80 degree Celsius until further processing.

### CD45 staining and FACS sorting

The cryopreserved leukocytic cells were briefly thawed at 37 degree Celsius and washed with complete RPMI 1640 (Cat. #10-040-CV, Corning) containing 10% FBS (Cat. #16140-071, gibco), 1% PSG (Cat. #P4333, Sigma), 10mM HEPES (Cat. #R5158, Sigma) and, incubated at 37 degree Celsius for 30 min. Post incubation, cells were washed with 2% FBS containing PBS and incubated with 1:20 FcR blocking reagent (Cat. #130-059-901, Miltenyi Biotec) in 2% PBS for 10 min at RT. The Fc blocked cells were stained with 1:200 diluted anti-human CD45-APC (Cat. #130-113-114, Miltenyi Biotec) for 20 min at 4 degree Celsius in dark. The CD45-stained cells were mixed with lived dead Sytox-Green stain (Cat. #S7020, ThermoFisher Scientific) and CD45^+^ cells were sorted with BD FACS Aria™III.

### Single Cell RNA Sequencing (scRNA-seq) Library Preparation

scRNA-seq was performed using 10x Genomics Chromium Single Cell Controller. Sorted cells were washed with 1X PBS and suspended in PBS/0.04% BSA. Cells were double-checked for viability and cell number by using the countess II FL and microscope. All cells were diluted to a concentration of 500-1000 cell/μl in PBS/0.04% BSA before being used for single cell 10X 3’v3. Single cells were captured using the 10X genomic controller according to the beads types and chip used for the experiments. The 10X genomic Chromium Single cell 3’ GEM, library and Gel bead Kit ‘v3 (cat. #1000075) and chromium chip B single cell kit (pat. #1000073) were used to capture cells on the controller; cell recovery targeted was in a range of 5000 - 10000 cells. Captured cells then undergo a GEM-RT, cDNA amplification, and all purification in accordance to the 10X protocol. Cleanup cDNA was checked via a tape station (Agilent 42000) HSD5000 (cat# part # 5067-5593) for cDNA traces. 25% of the cDNA was used to generate the library, and the Chromium i7 multiplex kit (part #120262) was used to identify each sample. Library cleanup was performed using AMPure beads and QC was done again with tape station D1000 tapes (Part # 5067-5583). Ten libraries of equal amount were pooled to give a final concentration of 10nM and submitted for sequencing with the NovaSeq6000 S2 sequencer, 28 cycles for read1, 8 cycles for i7 index, and 91 cycles for read 2 through the ATGC core at MD Anderson. Sequence data was then put through the 10X genomic cell ranger 3.0 pipeline. QC and Fastq files were obtained and checked for data quality, and Fastq files were used to do further analysis.

### RNA Isolation

RNeasy Mini Kit (Cat. #74104, Qiagen) was used to extract sorted cells and achieve efficient purification of total RNA from small amounts of starting material. The technology simplifies total RNA isolation by combining the stringency of guanidine-isothiocyanate lysis with the speed and purity of silica-membrane purification to ensure highest-quality RNA with minimum copurification of DNA. With the RNeasy Mini Kit, total RNA can be purified from 10 to 1 × 10^7^ animal or human cells, 0.5–30 mg human tissues. Briefly, samples are first lysed and then homogenized. Ethanol is added to the lysate to provide ideal binding conditions. The lysate is then loaded onto the RNeasy silica membrane. RNA binds (up to 100 μg capacity), and all contaminants are efficiently washed away. For certain RNA applications that are sensitive to very small amounts of DNA, the residual amounts of DNA remaining can be removed using a convenient on-column DNase treatment. Pure, concentrated RNA is eluted in 20–100 μl water.

### Low input mRNA sequencing

Illumina Compatible low input mRNA libraries were prepared using the Smart-Seq V4 Ultra Low Input RNA kit (Takara Bio, USA) and KAPA HyperPlus Library Preparation kit (Roche). Briefly, full length, double-stranded cDNA was generated from 8ng of total RNA using Takara’s SMART (Switching Mechanism at 5’ end of RNA Template) technology. The ds cDNA was amplified by nine cycles of LD-PCR, then purified using Ampure Beads (Agencort). Following bead elution, the cDNA was evaluated for size distribution and quantity using the Fragment Analyzer High Sensitivity NGS Fragment Analysis Kit (Agilent Technologies) and the Qubit dsDNA HS Assay Kit (ThermoFisher) respectively. The cDNA was enzymatically fragmented, and 20ng of the fragmented cDNA was used to generate Illumina compatible libraries using the KAPA HyperPlus Library Preparation kit. The KAPA libraries were purified and enriched with 2 cycles of PCR to create the final cDNA library. The libraries were quantified using the Qubit™ dsDNA HS Assay (ThermoFisher), then multiplexed 7 libraries per pool. The pooled libraries were quantified by qPCR using the KAPA Library Quantification Kit (KAPA Biosystems), and assessed for size distribution using the TapeStation 4200 (Agilent Technologies). The libraries were then sequenced, one pool per lane, on the Illumina HiSeq4000 sequencer using the 76bp paired end format.

### scRNA-seq Data Analysis

#### Raw sequencing data processing, quality check, data filtering, doublets removal, batch effect evaluation and data normalization

The raw scRNA-seq data were pre-processed (demultiplex cellular barcodes, read alignment, and generation of gene count matrix) using Cell Ranger Single Cell Software Suite provided by 10x Genomics. Detailed QC metrics were generated and evaluated. Cells with low complexity libraries or likely cellular debris (in which detected transcripts are aligned to less than 200 genes) were filtered out and excluded from subsequent analyses. Low-quality cells where >20% of transcripts derived from the mitochondria were considered apoptotic and also excluded. Following the initial clustering, likely cell doublets were removed from all clusters. Doublets were identified using a multi-step approach: 1) library complexity: cells with high complexity libraries (in which detected transcripts are aligned to more than 6500 genes) were removed; 2) Cluster distribution: doublets or multiplets likely form distinct clusters with hybrid expression features and exhibit an aberrantly high gene count; 3) cluster marker gene expression: cells of a cluster express markers from distinct lineages (e.g., cells in the T-cell cluster showed expression of myeloid cell markers and vice versa); 4) doublet detection algorithm: DoubletFinder^50^, an algorithm to predict doublets in scRNA-seq data, was applied to further identify and clean doublets that could have been missed by steps 1-3. We carefully reviewed canonical marker genes expression on UMAP plots and repeated the above steps multiple times to ensure elimination of most barcodes associated with cell doublets.

Following removal of poor-quality cells and doublets, a total of 144,678 cells were retained for downstream analysis. Library size normalization was performed in Seurat v3 (version 3.1.1)^51^ on the filtered gene-cell matrix to obtain the normalized UMI count as previously described^52^.

Statistical assessment of possible batch effects was performed using the R package k-BET (a robust and sensitive k-nearest neighbor batch-effect test) (PMID: 30573817). k-BET was run on cells from all samples, and on major cell types including microglia cells and CD8 T cells separately with default parameters. Each cell type was down sampled to 500 cells. We chose the k input value from 1% to 100% of the sample size. In each run, the number of tested neighborhoods was 10% of the sample size. The mean and maximal rejection rates were then calculated based on a total of 100 repeated k-BET runs. Following estimation of sample processing- or sequencing-related batch effects using k-BET, we employed Harmony for actual batch effect correction^53^. Harmony was run with default parameters to remove batch effects in the PCA space when clustering of major cell lineages before any clustering analysis or cell type identification/annotation was performed. We carefully evaluated the performance of Harmony in terms of its ability to integrate batches while maintaining cell type separation. Harmony was run on all cells to firstly identify major cell types. It was also run on each of the major cell types for subclustering analysis to further identify different cell states. To quantify the performance of Harmony, we further used k-BET and compared the rejection rate (reflecting batch effect) before and after Harmony. The data after Harmony showed a low rejection rate, indicating an excellent performance of batch effect correction in this study.

Moreover, we also applied the local inverse Simpson’s Index (LISI) to assess the performance of Harmony. As described previously^53^, the ‘integration LISI’ (iLISI) measures the degree of mixing among datasets (batches), ranging from 1 in an unmixed space to the number of datasets (batches) in a well-mixed space. And the ‘cell-type LISI’ (cLISI) measures integration accuracy using the same formulation but computed on cell-type labels instead. An accurate embedding has a cLISI close to 1 for every neighborhood, reflecting separation of different cell types. Before batch correction with Harmony, cells were mainly grouped by dataset (iLISI is around 1) and cells from different cell types were mixed (cLISI is far from 1). After batch correction with Harmony, iLISI and cLISI were re-computed in the Harmony embedding. iLISI is around 3.5, indicating a high degree of mixing among different datasets, and cLISI is very close to 1, reflecting excellent separation of different cell types while remain the well-mixed space.

#### Unsupervised cell clustering and dimensionality reduction

Seurat v3 (version 3.1.1)^51^ was applied to the normalized gene-cell matrix to identify highly variable genes (HVGs) for unsupervised cell clustering. To identify HVGs, the *vst* method in the Seurat package^51^ was used with default parameters. Principal component analysis (PCA) was performed on the top 2000 HVGs. The elbow plot was generated with the *ElbowPlot* function of Seurat^51^ and based on which, the number of significant principal components (PCs) were determined. The *FindNeighbors* function of Seurat was used to construct the Shared Nearest Neighbor (SNN) Graph, based on which the unsupervised clustering was done with *Seurat* function *FindClusters*. Different resolution parameters for unsupervised clustering were then examined in order to determine the optimal number of clusters. For visualization, the dimensionality was further reduced using Uniform Manifold Approximation and Projection (UMAP)^54^ method with *Seurat* function *RunUMAP*. The PCs used to calculate the embedding were as the same as those used for clustering. Two rounds of clustering were performed to identify major cell types (MG, non-MG myeloid cell, NK and T cells) and cell transcriptomic states within each major cell type. In the first round, 30-nearest neighbors of each cell were determined based on 30PCs to construct SNN graph. The clustering was performed with resolution 0.8 and each cluster was annotated by known markers (see ***Determination of major cell types and cell states***). Two major cell families comprising lymphoid- and, myeloid-cells, as well as negligible number of neural-like contamination were identified. The second-round clustering was performed on myeloid- and, lymphoid-cells respectively to identify cell states within each major cell types. For myeloid cell clustering, 40-nearest neighbors of each cell were determined based on 40 PCs to construct SNN graph. The clustering was performed with resolution 2, which resulted the identification of 34-cell states. For lymphoid cell clustering, the SNN graph was constructed based on 20-nearest neighbors of each cell that were determined by 30 PCs. The clustering was performed with resolution 2, which resulted the identification of 28-cell states.

#### Determination of major cell types and cell states

To define the major cell type of each single cell, differentially expressed genes (DEGs) were identified for each cell cluster using the *FindAllMarkers* analysis in the Seurat package^51^ and the top 50 most significant DEGs were carefully reviewed. The top 50 most significant differentially expressed transcription factors were also identified and reviewed by performing *FindAllMarkers* only on an aggregated TF list^55^. In parallel, feature plots were generated for top 20 DEGs and a suggested set of canonical lymphoid and myeloid cell markers, a similar approach as previously described^56,57^ followed by a careful manual review and annotation. The two approaches are combined to infer major cell types for each cell cluster according to the enrichment of marker genes and top-ranked DEGs in each cell cluster, and the global cluster distributions as previously described^57^. We sub-clustered the major myeloid and lymphoid populations iteratively and rigorously annotated the resulting cell-clusters using a combination of a) top 50 DEGs, b) top 50 differentially expressed lineage defining transcription factors and c) canonical immune signature genes.

#### Hierarchical relationship analysis

To study the hierarchical relationships among cell types identified in this study, unsupervised cluster analysis was performed. Pairwise Spearman correlations were calculated from average expression level (*Seurat* function *AverageExpression*) of each cell type, based on which Euclidean distances between cell types were calculated. Hierarchical cluster analysis was performed by R function *hclust* and the dendrogram was drawn using R package *ggtree*^58^.

#### Gene Ontology Enrichment Analysis

Top 100 DEGs from each cluster (MG and non-MG myeloid cells) were used for Gene Ontology enrichment analysis using Bioconductor package ClusterProfiler^59^. Significantly enriched GO-BP (Gene Ontology-Biological processes) terms were retrieved by setting the threshold of FDR<0.05 and minimum overlapping gene set size=3. Enriched terms having >=3 queried genes were manually selected using immunological keywords. Bubble plots were made using ggplot2 R package.

#### Macrophage Polarization Gene Set Enrichment Analysis

Top 100 DEGs from each cluster (MG, MAC and MDMs) were used for gene module enrichment analysis with reference gene signatures previously defined^27^ using Bioconductor package ClusterProfiler^59^ and plotted as circos plots^31^ using ggplot2 R package.

### Flow Cytometry staining

The cryopreserved cells were thawed at 37 degree Celsius and washed with10% FBS (Catalogue No. F4135, Sigma) in Iscove’s DMEM (1X, Catalogue No. 10-016-CV, Corning). The washed cells were pelleted by centrifugation at 500g for 5 mins. Cells were incubated with the 10% FBS containing Iscove’s DMEM media at 37°C for 30 mins before staining. PBS washed cells were stained with a fixable Live Dead Blue Dead cell stain dye (Catalogue No. L34962, Invitrogen) for 15 mins at 4°C. The Stained cells were washed with 1X PBS. Fc block was performed with a combination of Fc block -Human Tru Stain Fc block (Catalogue No. 422301, Biolegend), Nova Block Solution, (Catalogue No. M071437, Phitonex, Cell Blox Blocking Buffer, Catalogue No. B001T03F01) for 10 mins at 4°C. After Fc block, cell surface staining was performed with antibody cocktail (mentioned in **Supplementary Table S2)** diluted in BD Horizon Brilliant Stain buffer (Catalogue No. 566385, Becton Dickinson) and FACS buffer. For cell surface staining incubate was done for 30 mis at 4°C in dark. The stained cells after staining were washed with FACS buffer fixed with 200ul True Nuclear fixation buffer (True Nuclear 4X Fix Concentrate, Catalogue No. 73158, Biolegend and True Nuclear Fix Diluent, Catalogue No. 73160, Biolegend) overnight at 4°C. Overnight fixed cells were permeabilized with 1X Permeabilization buffer (True Nuclear 10X Perm, Catalogue No. 73162, Biolegend) for intracellular staining. Permeabilized cells were stained with the intracellular antibody cocktail (refer to **Supplementary Table S2**) for 20 mins at 4°C.The stained cells were resuspended in FACS buffer and data was acquired on Cytek Aurora 5 laser spectral flow cytometer.

### Cytometry Analyses

The acquired data was analyzed by Cytek SpectroFlo and Becton Dickinson FlowJo 10.8.1.

### Mouse experiments and survival analysis

Six-week-old WT and TREM2^-/-^ mice were obtained from Jackson laboratories. One week after guide screw implantation, 10,000 CT-2A cells were injected intracranially. Animals were monitored for tumor growth using bioluminescence imaging. For survival analysis, we used the log-rank test to calculate *P* values between groups, and the Kaplan-Meier method to plot survival curves using GraphPad Prism9 version 9.2.0 software.

---

**Supplementary Fig. S1.**
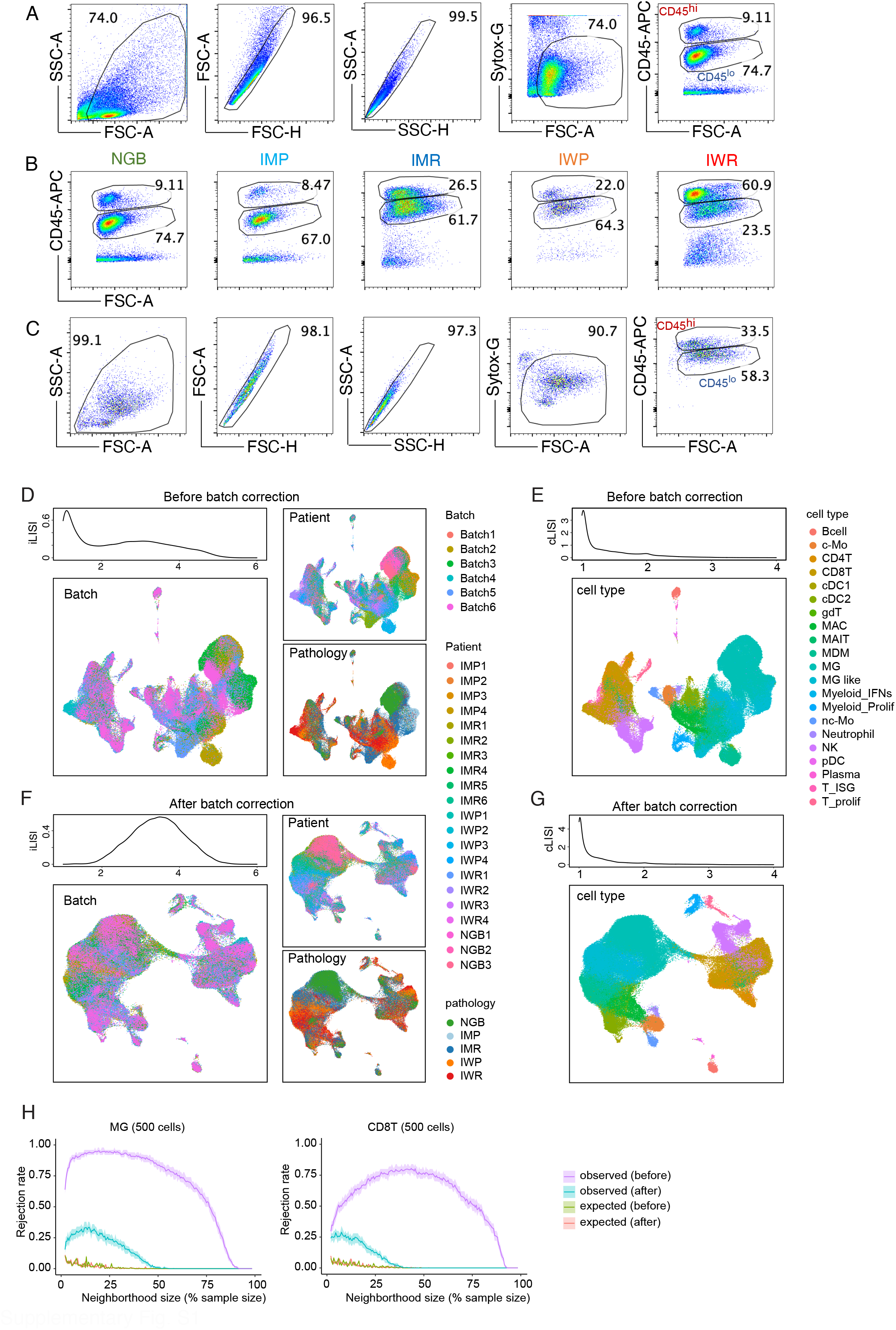
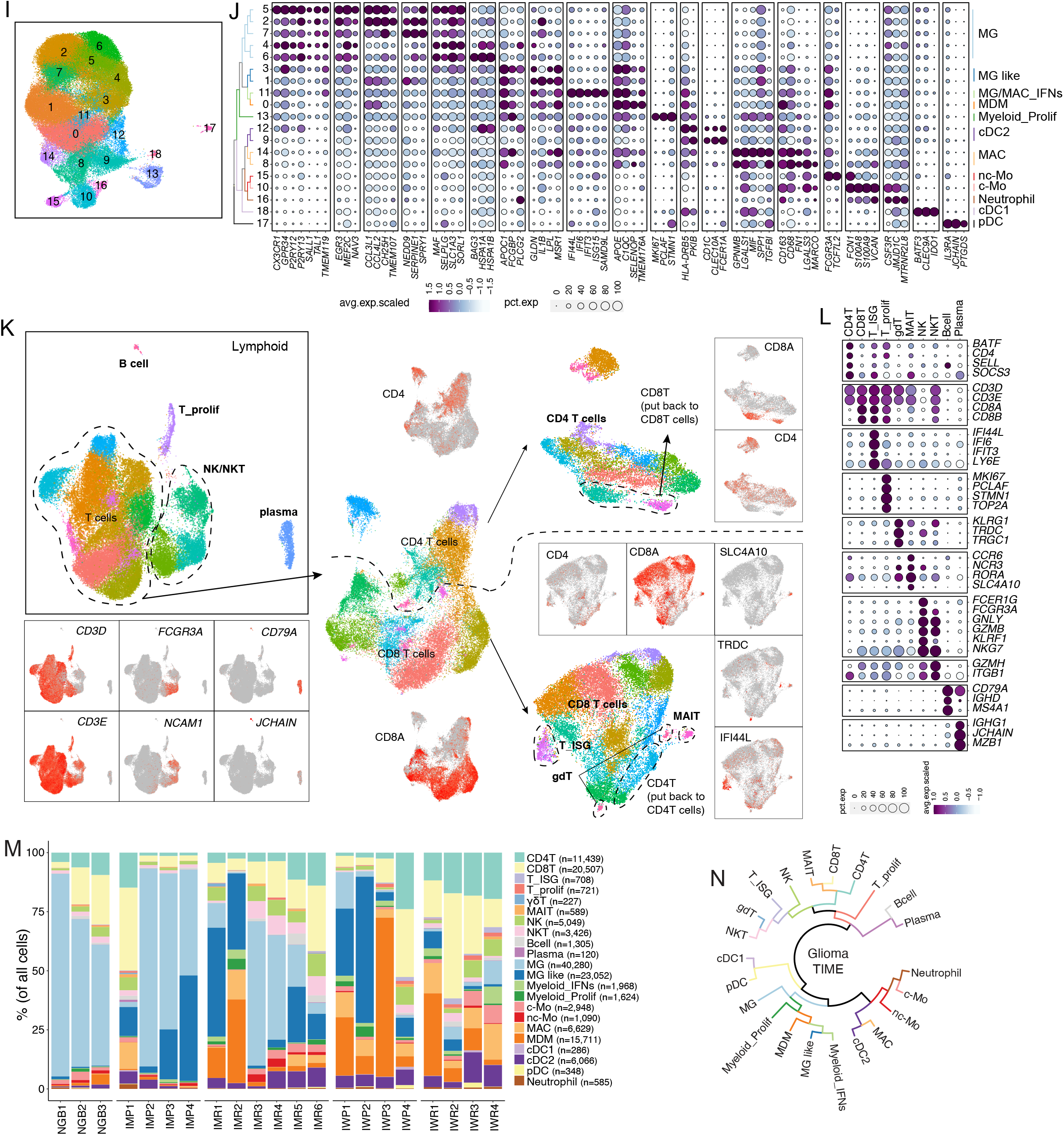
**(A):** Flow cytometry pseudo color dot plots showing in (A) gating strategy with SYTOX-G-stained dead cells and CD45 expression on stained tumor associated leukocytes, in **(B)** differential CD45 expression (CD45^lo^ and CD45^hi^) on leukocytes obtained from NGB controls and glioma subtypes as shown and in **(C)** representative pseudo color plots for purity of sorted CD45 cells. (**D**) iLISI (local inverse Simpson’s Index) computed for every cell’s neighborhood and summarized with density plots before batch correction. The UMAP were color coded by batch, patient and pathology. (**E**) cLISI computed for every cell’s neighborhood and summarized with density plots before batch correction. The UMAP were color coded by major cell types. (**F**) and **(G)**, same as **(D)** and **(E)** but after batch correction. **(H)** Batch effects evaluation using the k-nearest neighbor batch-effect test (k-BET, see Methods). K-BET was run on 500 cells randomly selected from 1 myeloid cell type (MG) and 1 lymphoid cell type (CD8T) separately before and after batch correction. Shaded areas represent the 95th percentile of n = 100 repeated k-BET runs. **(I)** UMAP visualization of unsupervised clustering analysis of glioma associated myeloid cells. **(J)** Bubble plot showing the scaled expression (shown by the color of the circle) and percentage of expression (shown by the size of the circle) of different myeloid cell type specific genes. **(K)** Workflow showing the identification of major lymphoid cell types by multi-steps of sub-clustering. **(L)** Bubble plot showing the scaled expression (shown by the color of the circle) and percentage of expression (shown by the size of the circle) of different lymphoid cell type specific genes. **(M)** The stacked bar plot showing relative distribution of glioma associated myeloid and lymphoid cell types across sc-RNAseq sampled patients. **(N)** Phylogenic relationship between myeloid and lymphoid cell types revealed by hierarchical clustering analysis based on Euclidean distance between cell types (see Methods).

**Supplementary Fig. S2.**
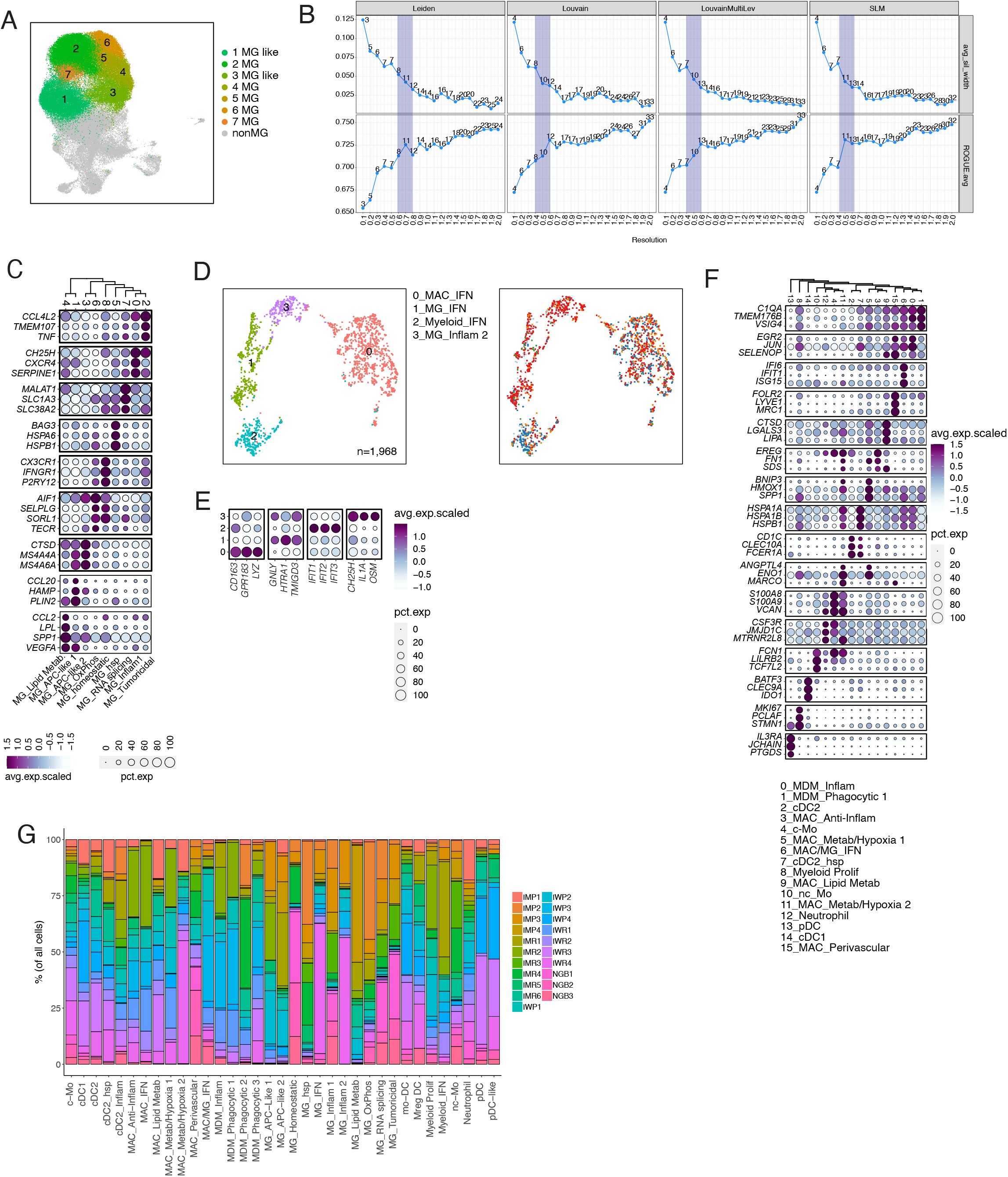
**(A):** UMAP visualization of unsupervised clustering analysis of glioma associated MG, MG-like (colored) and non-MG myeloid (grey). **(B)** Four different community detection algorithms in Seurat were used to find the best clustering of MG cells. In each algorithm, resolution was tuned from 0.1 to 2 with step 0.1, under which the averaged silhouette width value and ROGUE value were calculated to evaluate the inter-cluster dissimilarity and cluster purity. Shaded region indicated the best trade-off between averaged silhouette width value and ROGUE value. **(C)** Bubble plot showing the scaled expression (shown by the color of the circle) and percentage of expression (shown by the size of the circle) of different lymphoid cell type specific genes. **(D)** UMAP visualization of unsupervised clustering analysis of glioma associated Myeloid interferons (Myeloid_IFN). Cells are color coded for their inferred cell types (left) and the glioma subtypes of their corresponding tumors (right). **(E-F)** Bubble plot showing the scaled expression (shown by the color of the circle) and percentage of expression (shown by the size of the circle) of different Myeloid interferon clusters and their specific genes in (E) and non-MG myeloid cells in (F). **(G)** Stacked bar plots showing corresponding %myeloid cell type composition across each glioma subtype.

